# Heterochromatin dynamics during the initial stages of sexual development in *Plasmodium falciparum*

**DOI:** 10.1101/2024.03.19.585770

**Authors:** Sandra Nhim, Elisabet Tintó-Font, Núria Casas-Vila, Lucas Michel-Todó, Alfred Cortés

## Abstract

Asexual replication of *Plasmodium falciparum* in the human blood results in exponential parasite growth and causes all clinical symptoms of malaria. However, at each round of the replicative cycle, some parasites convert into sexual precursors called gametocytes, which are essential for transmission to mosquito vectors. After sexual conversion, parasites develop through the sexual ring stage and then gametocyte stages I to V before they are infective to mosquitoes. Heterochromatin, a type of chromatin generally refractory to gene expression, plays an important role in the regulation of sexual conversion by silencing the master regulator of the process, PfAP2-G, in asexual parasites. Additionally, previous reports have described changes in the genome-wide distribution of heterochromatin in stage II/III or older gametocytes, including expansion of heterochromatin at several subtelomeric regions and reduced occupancy at a few specific loci. However, it is not known if these changes occur concomitantly with sexual conversion or at a later time during gametocyte development. Using a transgenic line in which sexual conversion can be conditionally induced, here we show that the genome-wide distribution of heterochromatin at the initial stages of sexual development (i.e., sexual rings and stage I gametocytes) is almost identical to parasites at asexual blood stages, and major changes do not occur until stage II/III. We also show that transcriptional changes associated with sexual development typically precede, rather than follow, changes in heterochromatin occupancy at their loci, which raises the possibility that PfAP2-G operates as a pioneer factor.

**IMPORTANCE:** Epigenetic processes and chromatin structure play an important role in the regulation of gene expression in malaria parasites. In particular, a type of chromatin called heterochromatin is involved in the regulation (silencing) of many genes. Parasite sexual development is essential for transmission to mosquito vectors. Here we characterised the global distribution of heterochromatin at different stages of sexual development, and found that initially it is identical to asexual blood stages, but at later transmission stages it is altered. This informs about the putative roles of general heterochromatin redistribution in parasite life cycle progression. By integrating multi-omic datasets, we also found that changes in the expression of several genes precede changes in their heterochromatin occupancy. This indicates that during sexual development some genes can be activated in spite of having heterochromatin.

## INTRODUCTION

In eukaryotic organisms, chromatin plays fundamental roles in the regulation of gene expression and other nuclear processes. A type of chromatin termed heterochromatin mediates transcriptional repression and also plays important roles in centromere and telomere function and in repressing transposable elements, thus securing chromosome stability [1–4]. Conserved features of heterochromatin across eukaryotes include its ability to spread into adjacent regions and epigenetic inheritance of its distribution during cell division. Regions that are found as heterochromatin in some cells and as euchromatin in others, which often include protein-coding genes, are referred to as facultative heterochromatin, whereas regions that are always found as heterochromatin, such as subtelomeric and pericentromeric tandem repeats regions, are called constitutive heterochromatin. In the majority of eukaryotes, the main molecular determinants of heterochromatin are the histone modifications di- or tri-methylation of histone H3 lysine 9 (H3K9me2 or H3K9me3) or tri-methylation of lysine 27 (H3K27me3). H3K27me3 is typically associated with dynamic facultative heterochromatin and regulates transient gene silencing during development and cell type specification in multicellular organisms. In contrast, H3K9me3 is the hallmark of constitutive heterochromatin, although it can also play important roles in the dynamic regulation of gene expression by forming facultative heterochromatin [1–4].

In *Plasmodium falciparum*, the parasite responsible for the most severe forms of human malaria, H3K9me3 is dynamic and frequently associated with facultative heterochromatin, whereas H3K27me3 has been identified only at specific stages of sexual development [5] and its function still remains unclear. H3K9me3 has been extensively studied, as it plays fundamental roles in parasite biology. This mark, together with the conserved heterochromatin protein 1 (HP1) that binds to it, occupies subtelomeric repeats and clonally variant gene (CVG) loci, which are located in subtelomeric regions and a few chromosome internal islands [6–10]. Presence of H3K9me3 and HP1 at the upstream regulatory regions of CVGs, where they form facultative heterochromatin, is associated with transcriptional silencing, whereas presence of acetylated H3K9 (H3K9ac) at these positions is associated with their active state [9]. The distribution of H3K9me3-based heterochromatin at CVGs is clonally transmitted from one generation of blood stage asexual parasites to the next, acting as a truly epigenetic mark [11]. However, infrequent transitions between the heterochromatic and euchromatic states result in transcriptional switches [12–14].

The *P. falciparum* genome contains more than 500 CVGs, which participate in numerous host-parasite interactions, including antigenic variation, solute transport, erythrocyte invasion, erythrocyte remodelling and sexual conversion [12–15]. Many of these genes belong to multigene families in which there is redundancy, as different genes of the same family often encode similar proteins that participate in the same process, albeit with antigenic or functional differences. Therefore, switches in the expression of CVGs ultimately result in antigenic or phenotypic variation. In general, epigenetic regulation of malarial CVGs plays an adaptive role: functional or antigenic diversity within parasite populations provides the grounds for dynamic natural selection when the conditions of the environment change. This is considered a bet-hedging adaptive strategy [15].

The *P. falciparum* life cycle involves multiple well-differentiated stages in humans and in *Anopheles* spp. mosquito vectors. In the human blood, parasites undergo the asexual intraerythrocytic development cycle (IDC), which lasts ∼48 h and involves the merozoite, ring, trophozoite and schizont stages. Repeated rounds of the IDC result in exponential parasite growth, which is associated with all clinical symptoms of malaria. However, at each round of the IDC, a small fraction of the parasites abandons asexual growth and converts into sexual precursors called gametocytes, in a process called sexual conversion. Gametocytes are the only form of the parasite that, once mature, can infect mosquitoes. In the mosquito, after mating of male and female gametocytes and several additional stage transitions, parasites become salivary gland sporozoites, which can infect a new human host during a mosquito bite. Sporozoites invade hepatocytes, where they multiply until they are released to the blood stream to start a new blood infection, closing the cycle [16].

The consolidated model for sexual conversion postulates that PfAP2-G, an ApiAP2 transcription factor, is the master regulator of the process [17–20]. During the IDC, the *pfap2-g* locus is in a heterochromatic state that maintains the gene silenced. Expression of the GDV1 protein mediates heterochromatin depletion at the *pfap2-g* locus [21], which results in expression of PfAP2-G and sexual conversion. The heterochromatin-based regulation of PfAP2-G [17, 22] illustrates the cross-talk between transcription factor networks and epigenetic regulation of gene expression. Heterochromatin may also play a role in the regulation of *gdv1*, as heterochromatin occupancy at this locus correlates with basal sexual conversion rates [9]. PfAP2-G regulates the expression of several early gametocyte genes, which drive sexual differentiation [20]. Depending on the time of activation of PfAP2-G expression during the IDC, parasites can convert directly into gametocytes via the same cycle conversion (SCC) pathway, or go through one additional cycle of multiplication as sexually-committed forms before converting via the next cycle conversion (NCC) pathway [23]. Regardless of the conversion pathway used, a cascade of transcription factors activation ensues [24, 25] and drives development through the sexual ring stage and then stage I to V gametocytes, in a process that lasts ∼10 days until male or female gametocytes are mature and ready to infect a mosquito.

Heterochromatin distribution has been compared between clonal parasite lines that differ in the expression of specific CVGs, which revealed alternative heterochromatin patterns associated with the active or silenced states of these genes [9, 12, 26–28]. However, heterochromatin distribution differences associated with life cycle progression have also been reported. While several studies found that the distribution of heterochromatin at specific *P. falciparum* CVG loci [26–28] or at a genome-wide level [7] is almost identical between different stages of the IDC, the genome-wide distribution of heterochromatin is different in gametocytes [7, 29] or mosquito stages (oocysts and sporozoites) [30, 31]. Heterochromatin distribution is also different between asexual blood stages and sporozoites in the simian malaria parasite *P. cynomolgi* [32], which suggest that heterochromatin remodelling during transmission stages is a conserved feature of *Plasmodium* spp. The most prominent heterochromatin alterations observed during transmission stages were expansion of heterochromatic domains at some subtelomeric regions and opening of heterochromatin at a small number of specific loci. Another recent study reported heterochromatin differences between male and female gametocytes [33]. However, none of these studies included early sexual stages such as sexual rings or stage I gametocytes. Therefore, it is currently not known if changes in heterochromatin distribution during sexual development occur concomitantly with sexual commitment, at the initial stages of sexual development or later on during sexual development.

Here we took advantage of a recently developed inducible parasite line that enables controlled massive sexual conversion [19] to investigate heterochromatin distribution in early sexual stages. To assess the impact of heterochromatin redistribution during sexual development on gene expression, we characterised the relative temporal dynamics of changes in heterochromatin and in gene expression.

## RESULTS

### Heterochromatin distribution is almost identical between asexual blood stages, sexual rings and stage I gametocytes

To characterise the distribution of heterochromatin at the initial stages of *P. falciparum* gametocyte development, we used the transgenic line E5ind, in which synchronous sexual conversion of the vast majority of parasites can be conditionally induced [19]. In brief, addition of rapamycin to E5ind cultures at the trophozoite stage results in activation of the expression of PfAP2-G and sexual conversion via the NCC pathway [23] in ∼90% of the parasites, such that they develop into sexually committed schizonts and, after reinvasion, into sexual rings and subsequent stages of sexual development (stage I to V gametocytes) [19].

Using the E5ind line, we performed H3K9me3 chromatin immunoprecipitation followed by sequencing (ChIP-seq) for sexual rings, stage I and stage II/III gametocytes, and also for asexual rings prepared in parallel without adding rapamycin (Fig. 1A and Supplementary Fig. S1). Since heterochromatin distribution remains stable throughout the full IDC [7], the asexual ring stage is representative of all other IDC stages. Visual inspection of the general distribution of heterochromatin in two independent biological replicates did not reveal any major differences between asexual rings, sexual rings and stage I gametocytes, whereas in stage II/III gametocytes expansion of heterochromatin was apparent at several subtelomeric regions (Fig. 1B). Pearson correlation analysis confirmed a very high level of heterochromatin distribution similarity in pairwise comparisons between different stages (*r* > 0.93 in all comparisons), with the two replicates of stage II/III gametocytes forming a separate cluster (Sup. Fig. S2A). Consistently, the proportion of the genome covered by a H3K9me3 peak (according to MACS2 peak calling) was slightly higher in stage II/III gametocytes than at the other stages (Sup. Fig. S2B). These results indicate that heterochromatin distribution is almost identical between asexual blood stages and sexual rings or stage I gametocytes, and that the previously reported changes in heterochromatin distribution during sexual development [7, 29] do not occur until at least the stage II of gametocyte development.

**Fig. 1.**
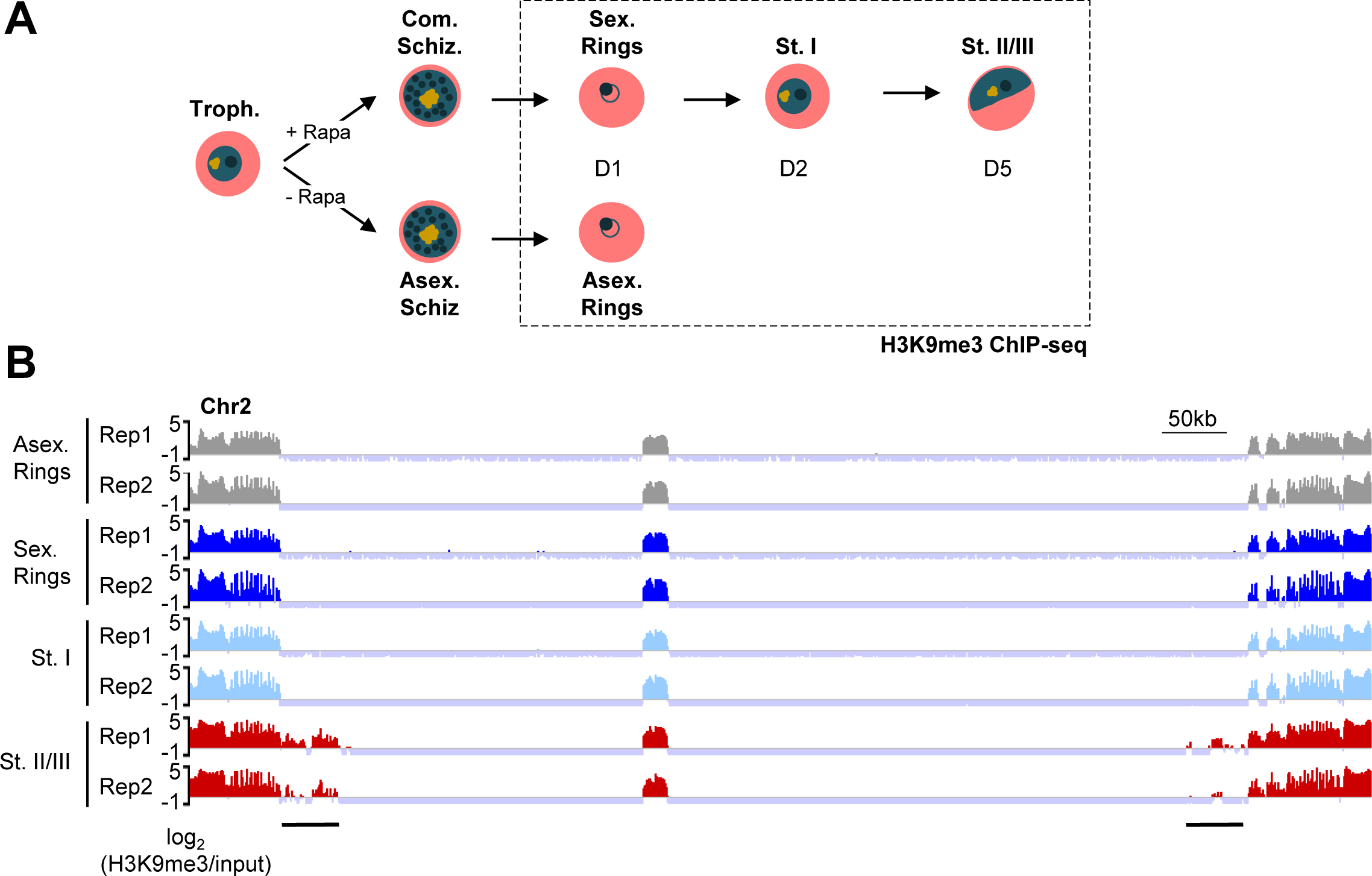
Overview of heterochromatin dynamics at the early stages of sexual development. **A.** Schematic of the experiment design. Sorbitol-synchronised E5ind cultures at the trophozoite (Troph.) stage were treated with rapamycin (+ Rapa) to induce sexual conversion, or with DMSO solvent (- Rapa). Chromatin was extracted for ChIP-seq analysis on the days (D) post-induction indicated. The stages analysed were asexual (Asex.) rings (D1), sexual (Sex.) rings (D1), stage I gametocytes (St. I, D2) and stage II/III gametocytes (St. II/III, D5). Committed Schizonts (Com. Schiz.) and Asexual Schizonts (Asex. Schiz.) were not analysed. **B.** H3K9me3 distribution in the full chromosome 2 (as a representative example) at different asexual and sexual blood stages. Replicates (Rep) 1 and 2 are biological replicates. Black horizontal lines indicate the heterochromatin expansions observed in stage II/III gametocytes.

### Subtelomeric heterochromatin expansions occurring from gametocyte stage II/III are similar among parasite lines of different genetic backgrounds

Since the most prominent heterochromatin distribution change during sexual development was expansion of heterochromatin at several subtelomeric regions, we developed an in-house pipeline to measure the length of the region covered by heterochromatin at each of the 28 subtelomeric regions and also at seven internal heterochromatin islands. The method was based on MACS2 peak calling and joining peaks within the same subtelomeric regions or internal island (Sup. Fig. S3A). This analysis confirmed that the size of all heterochromatic regions was almost identical between asexual parasites, sexual rings and stage I gametocytes, whereas marked heterochromatin expansions (relative to asexual parasites) occurred in several subtelomeric regions in stage II/III gametocytes (Fig. 2A and Sup. Fig. S3B). Heterochromatin islands remained invariant across all stages (Sup. Fig. S4).

**Fig. 2.**
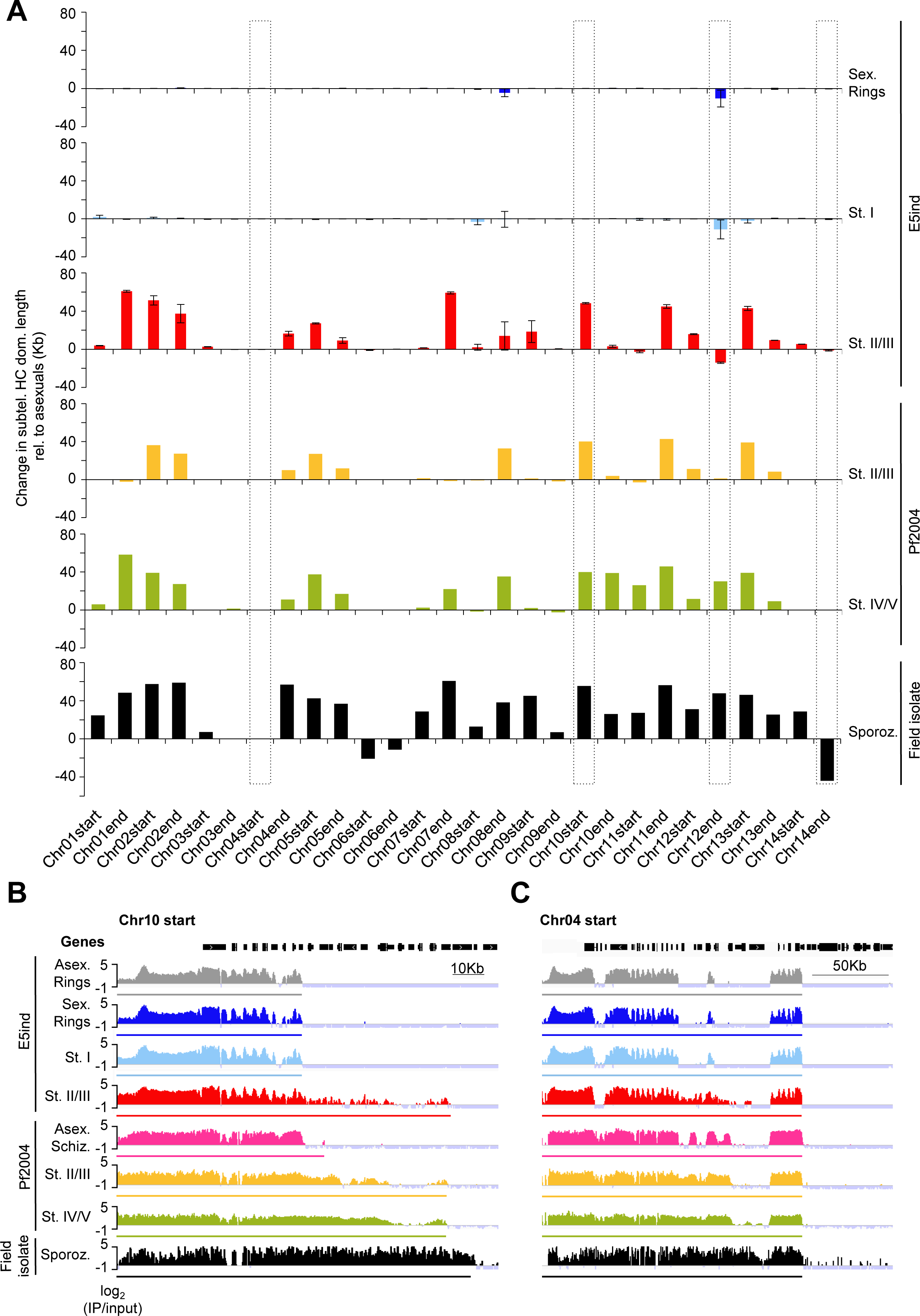
Changes in the extension of heterochromatin domains during sexual development. **A.** Changes in the size of subtelomeric heterochromatin (HC) domains at different stages of sexual development. Values for the E5ind line (H3K9me3 ChIP-seq, this study) are changes relative to E5ind asexual parasites (asexual rings) and are the average of two independent biological replicates, with S.E.M. An analogous analysis of data from two previously published studies is included for comparison. Values for the Pf2004 line (HP1 ChIP-seq, single replicate) [7] are changes relative to Pf2004 asexual parasites (schizonts). Values for the Field isolate sporozoites (H3K9me3 ChIP-seq, single replicate) [30] are changes relative to E5ind asexual rings. Dotted boxes indicate regions discussed in the text. Abbreviations for parasite stages are as in Fig. 1. **B-C.** Representative examples of a subtelomeric heterochromatin expansion observed from stage II/III gametocytes onwards (**B**) and a heterochromatin expansion that does not affect the limits of the subtelomeric heterochromatin domain (**C**). The coloured lines below each ChIP-seq track indicate the bioinformatically estimated extension of subtelomeric heterochromatin domains. The ChIP-seq values for the E5ind line are the mean of replicates 1 and 2.

Next, we compared the size of subtelomeric or internal heterochromatin domains between our samples and published heterochromatin (H3K9me3 or HP1) ChIP-seq data from stage II/III or IV/V gametocytes of the Pf2004 genetic background [7], and sporozoites from a field isolate [30]. Overall, similar heterochromatin expansions (relative to asexual parasites), affecting the same subtelomeric regions and with similar limits, were observed between our E5ind stage II/III gametocytes and Pf2004 gametocytes [7] (Fig. 2A-B and Sup. Fig. S4-7). The few different expansions between E5ind and Pf2004 could be explained by differences in heterochromatin distribution between the two lines that were already present in asexual parasites, as a consequence of regular epigenetic variation [9, 15] (i.e., some genes are silenced and heterochromatic in asexual parasites in one parasite line and active and euchromatic in the other). However, in many of the regions where the expansions occurred, H3K9me3 coverage was low compared with other heterochromatic regions, mainly in our E5ind stage II/III gametocytes (Fig. 1B, 2B and Sup. Fig. S3A). This is suggestive of population heterogeneity, such that heterochromatin had already expanded in older individual gametocytes but not yet in the younger ones. Of note, heterochromatin distribution in sporozoites [30] was generally similar to stage II/III or IV/V gametocytes (Fig. 2A-B and Sup. Fig. S5B), although some additional expansions were observed. Since ChIP-seq data from asexual parasites of the same genetic background as the sporozoites sample was not available, changes in heterochromatin distribution in sporozoites had to be determined relative to E5ind asexual parasites. Therefore, some of the differences in heterochromatin expansions between gametocytes and sporozoites may be attributable to the different genetic backgrounds.

In addition to heterochromatin expansions, an apparent retraction of heterochromatin was observed at the distal end of chromosome 12 in E5ind stage II/III gametocytes and at the distal end of chromosome 14 in sporozoites (Fig. 2A), but this could be explained by small changes at a specific locus or by loss of heterochromatin in two genes expressed in mosquito stages, respectively (Sup. Fig. S6A). In addition to the changes in the limits of specific subtelomeric heterochromatin domains, there were changes in heterochromatin distribution within some of the domains. In some subtelomeric regions there was an expansion of heterochromatin that did not affect the estimated limit of the domain, involving loci that were euchromatic in asexual parasites (Fig. 2C).

Together, these results show that during sexual development there is expansion of heterochromatin at specific subtelomeric regions, starting at the gametocyte stage II or III. The heterochromatin expansions observed were similar between parasites of different genetic background, indicating that this is a conserved, controlled process intrinsic to *P. falciparum* sexual development.

### Distribution of specific changes in heterochromatin coverage during sexual development

To analyse in more detail the changes in heterochromatin distribution, regardless of whether or not they affect the size and limits of large heterochromatin domains, we modified a previously described custom differential peak calling method [9] to identify regions with differential H3K9me3 coverage between different stages. Comparison of E5ind asexual stages and stage II/III gametocytes revealed 139 differential coverage regions, of which 33.8% were localised in subtelomeric repeats regions devoid of coding genes and containing mainly TARE repeats [34] (Fig. 3A and Sup. Table S1). Heterochromatin coverage in these subtelomeric repeats was generally higher in sexual compared to asexual parasites. Of note, 21 of the previously reported 22 subtelomeric lncRNA-TARE transcripts [35] intersected with differential coverage regions (Sup. Fig. S8 and Sup. Table S1).

**Fig. 3.**
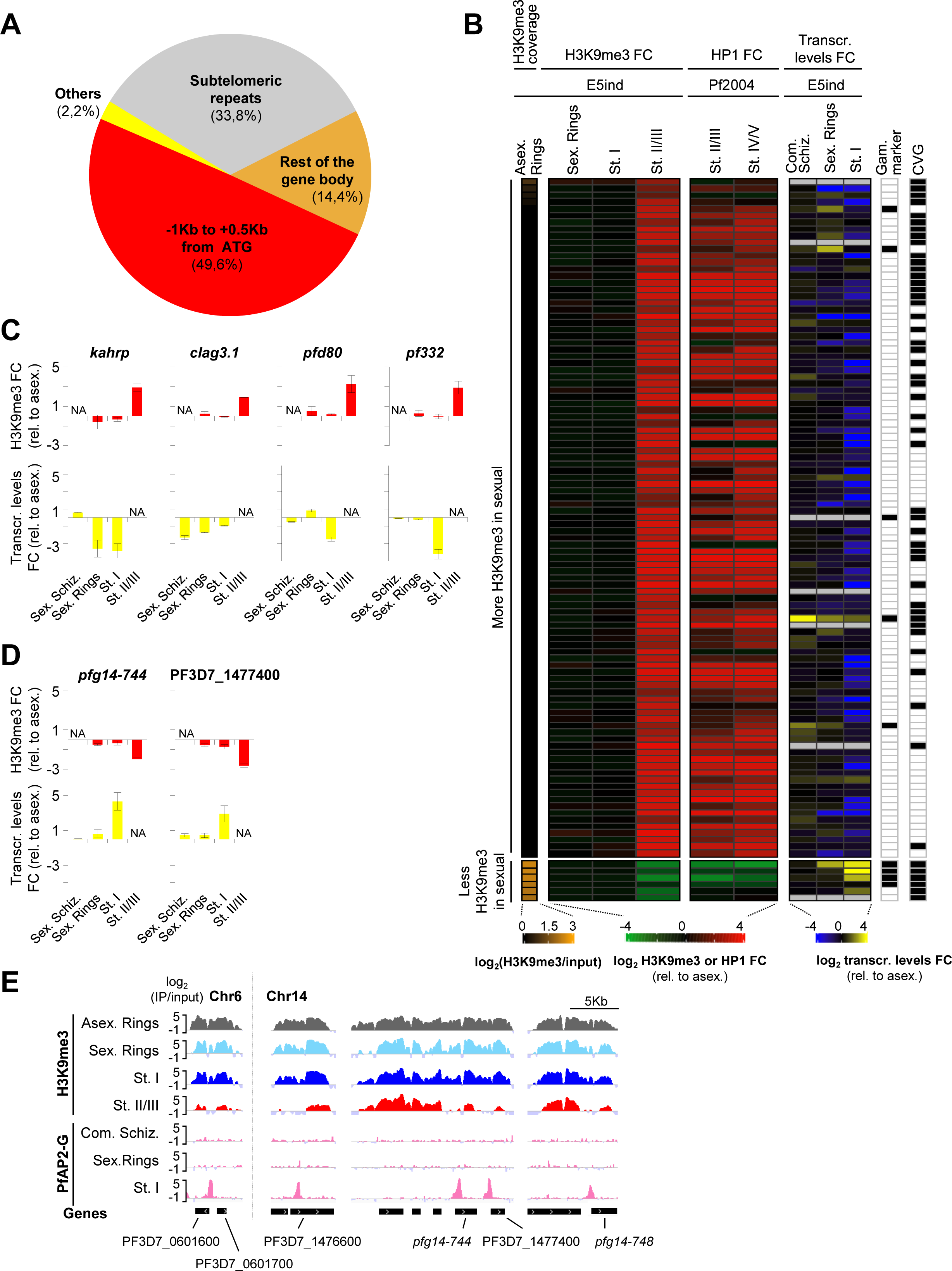
Genes with differential heterochromatin coverage during sexual development. **A.** Characteristics of 139 regions with different H3K9me3 coverage between E5ind asexual rings and stage II/III gametocytes. **B.** Heatmap showing changes in heterochromatin coverage and transcript levels during sexual development for 107 stage II/III differential coverage genes. Coverage values were calculated at the region −1,000 to +500 bp from the ATG. Values in the first column are H3K9me3 coverage in E5ind asexual parasites, whereas the following columns are coverage fold-change (FC) at different sexual stages relative to asexual parasites. Published HP1 ChIP-seq coverage FC for Pf2004 gametocytes (relative to asexual parasites) [7] is shown for comparison. The next columns are published E5ind transcript levels FC at different sexual stages relative to their asexual counterparts [19]. Values are the average of two independent biological replicates (E5ind ChIP-seq and transcript levels) or the result of a single experiment (Pf2004). Grey indicates that no data was available. Genes previously classified as gametocyte markers or as CVGs, according to previously published lists [9, 36], are indicated. Abbreviations of stages are as in Fig. 1. **C-D.** Log_2_ of the H3K9me3 coverage FC relative to asexual parasites (top) and transcript levels FC relative to asexual counterparts [19] (bottom) at different stages of sexual development. Representative genes with increased (**C**) or reduced (**D**) H3K9me3 coverage in stage II/III gametocytes are shown. Values are the average of two biological replicates, with S.E.M. N.A.: not analysed. **E.** PfAP2-G binding at loci in which heterochromatin coverage is reduced in E5ind stage II/III gametocytes. AP2-G data is from a published study using the E5-derived AP2-G-DD line (average of two independent biological replicates) [20]. Arrows within genes indicate the direction of transcription. E5ind H3K9me3 values are the average of two independent biological replicates.

A large proportion (49.6%) of the 139 differential peaks overlapped with the −1,000 to +500 bp (from the ATG) region of a gene, and 14.4% of the peaks overlapped only with other parts of a gene (including the 3’UTR, when annotated). Lastly, three of the differential peaks (2.2%) occurred in intergenic regions that did not overlap with any of the annotated genetic elements mentioned above (“Others”) (Fig. 3A).

In contrast to the 139 differential H3K9me3 coverage regions identified when comparing asexual rings with stage II/III gametocytes, only 21 and two differential coverage regions were identified between asexual rings and sexual rings or between asexual rings and stage I gametocytes, respectively (Sup. Table S1). In the case of sexual rings, the majority (76%) of the differential coverage regions overlapped with the −1,000 to +500 bp region of a gene (Sup. Fig S9A), whereas the two regions detected in stage I gametocytes were intergenic.

### Genes with differential heterochromatin coverage at putative regulatory regions between asexual and sexual parasites

Given that heterochromatin changes at proximal upstream regions and the beginning of the coding sequence show the strongest association with gene expression [9], we analysed in detail the differential heterochromatin coverage regions overlapping with the −1,000 to + 500 bp (from the ATG) region of a gene. In the comparison between asexual rings and stage II/III gametocytes, 107 genes were identified in which this region overlapped with a differential H3K9me3 peak and had a coverage difference higher than twofold (Figure 3B and Sup. Table S2). These will be referred to as ‘differential coverage genes’. For these genes, H3K9me3 coverage in the −1,000 to + 500 bp region was almost identical between E5ind asexual rings, sexual rings and stage I gametocytes, and markedly different only in stage II/III gametocytes. Heterochromatin changes at these genes in stage II/III gametocytes were similar to changes in stage II/III or IV/V gametocytes of the Pf2004 line [7] (Figure 3B). Indeed, there was a large overlap between the lists of differential heterochromatin coverage genes in E5ind stage II/III and Pf2004 gametocytes (Sup. Table S3).

101 of the E5ind stage II/III differential coverage genes showed an increase in heterochromatin occupancy in stage II/III gametocytes compared to asexual parasites, reflecting mainly the previously described subtelomeric heterochromatin expansions. The majority of these 101 genes were not gametocyte markers and about half were previously described as CVGs [36], with a predominance of genes encoding exported proteins involved in erythrocyte remodelling and PfEMP1 export, consistent with previous reports [7] (Figure 3B and Sup. Table S2). However, in the remaining six differential coverage genes there was a reduction of heterochromatin coverage. For these genes, reduced occupancy was also observed only in stage II/III gametocytes. These genes included a cluster of early gametocyte markers located in the distal subtelomeric region of chromosome 14, including *pfg14-744* and *pfg14-748* [37], and two neighbour genes in chromosome 6 (Figure 3B and Sup. Table S2).

In contrast to the many differential coverage genes in stage II/III gametocytes (relative to asexual rings), an analogous analysis for sexual rings and stage I gametocytes revealed only seven and zero differential coverage genes, respectively (Sup. Fig. S9B-C and Sup. Table S4). In all seven differential coverage genes in sexual rings, there was a decrease in coverage of very low magnitude. Furthermore, the decrease was no longer observed in stage I or II/III gametocytes, suggesting that differential coverage in these few genes was likely explained by technical reasons.

To investigate the transcriptional changes during sexual development associated with the changes in heterochromatin occupancy, we used published gene expression data for the E5ind line [19]. This analysis included the sexually-committed schizont, sexual ring and stage I gametocyte stages and their asexual counterparts (asexual schizont, asexual ring and trophozoite). In general, there were lower transcript levels in sexual stages (compared with the asexual counterparts) for the genes that gained heterochromatin during sexual development, and higher transcript levels for genes that lost heterochromatin. However, for many of the genes, transcript level changes already occurred in stage I gametocytes or even earlier sexual stages, in contrast to heterochromatin changes that were not observed until stage II/III gametocytes (Fig. 3B).

### Transcriptional changes during sexual development may precede heterochromatin changes

To explore in more detail the relative timing of transcriptional and heterochromatin changes, we compared the temporal dynamics of heterochromatin coverage and transcript levels during sexual development for selected representative genes (using the same data as in Fig. 3B). For these genes, changes in heterochromatin did not occur until stage II/III gametocytes, whereas changes in transcript levels [23] occurred at an earlier stage (Figure 3C-D). In the genes *kahrp*, *clag3.1*, *pfd80* or *pf332*, increased H3K9me3 coverage was not observed until stage II/III gametocytes, whereas reduced transcript levels (compared with their asexual counterparts) were already observed at the sexually-committed schizont (*clag3.1*), sexual ring (*kahrp*) or stage I gametocyte (*pfd80* and *pf332*) stages (Fig. 3C). Likewise, reduced H3K9me3 coverage was observed in *pfg14-744* and a neighbour *phist* gene (PF3D7_1477400) only in stage II/III gametocytes, but increased transcript levels were already observed in stage I gametocytes (Figure 3D). All six genes with reduced heterochromatin occupancy during sexual development are direct targets of PfAP2-G [20]. Comparison with published PfAP2-G binding data (ChIP-seq analysis) [20] revealed that this transcription factor already binds to the putative regulatory region of these genes in stage I gametocytes, at positions that overlap with the regions where heterochromatin occupancy is reduced (Figure 3E). Therefore, the temporal dynamics of PfAP2-G binding appears to correlate with changes in gene expression and precede changes in heterochromatin coverage.

To confirm these results, we performed RT-qPCR and H3K9me3 ChIP-qPCR analysis from time-course experiments that included five time points from the sexual ring to stage III gametocyte stages, with a non-induced asexual rings culture as a control (Fig. 4A). In addition to genes in Fig. 3C-D, in this analysis we also included *pfg14-748,* PF3D7_0601600 (putative tetratricopeptide repeat protein), for which data from the previous microarray-based transcriptomic analysis [19] was not available, and *geco*, which did not pass the threshold for differential heterochromatin coverage but showed a peculiar pattern of early heterochromatin alterations (Sup. Fig. S6A). For all genes analysed, the same trend (increase or decrease) in heterochromatin occupancy during sexual development as in the ChIP-seq analysis was observed. These experiments confirmed that changes in H3K9me3 coverage generally occurred from stage II or II/III onwards, whereas transcript levels started to increase or decrease one to two days earlier (Fig. 4B-C). In general, in genes with increased H3K9me3 coverage the change was observed from day 5 post-induction (stage II/III gametocytes), whereas in genes with reduced coverage it was observed from day 3 (stage II gametocytes). Changes in transcript levels during sexual development (silencing of genes with increased heterochromatin or activation of genes with reduced heterochromatin) were typically observed from day 2 (stage I gametocytes) or even earlier, confirming that these transcriptional changes precede, rather than follow, heterochromatin changes. For *pf14-744* and *pf14-748*, high transcript levels occurred transiently only until stage II gametocytes, as previously reported [37, 38].

**Fig. 4.**
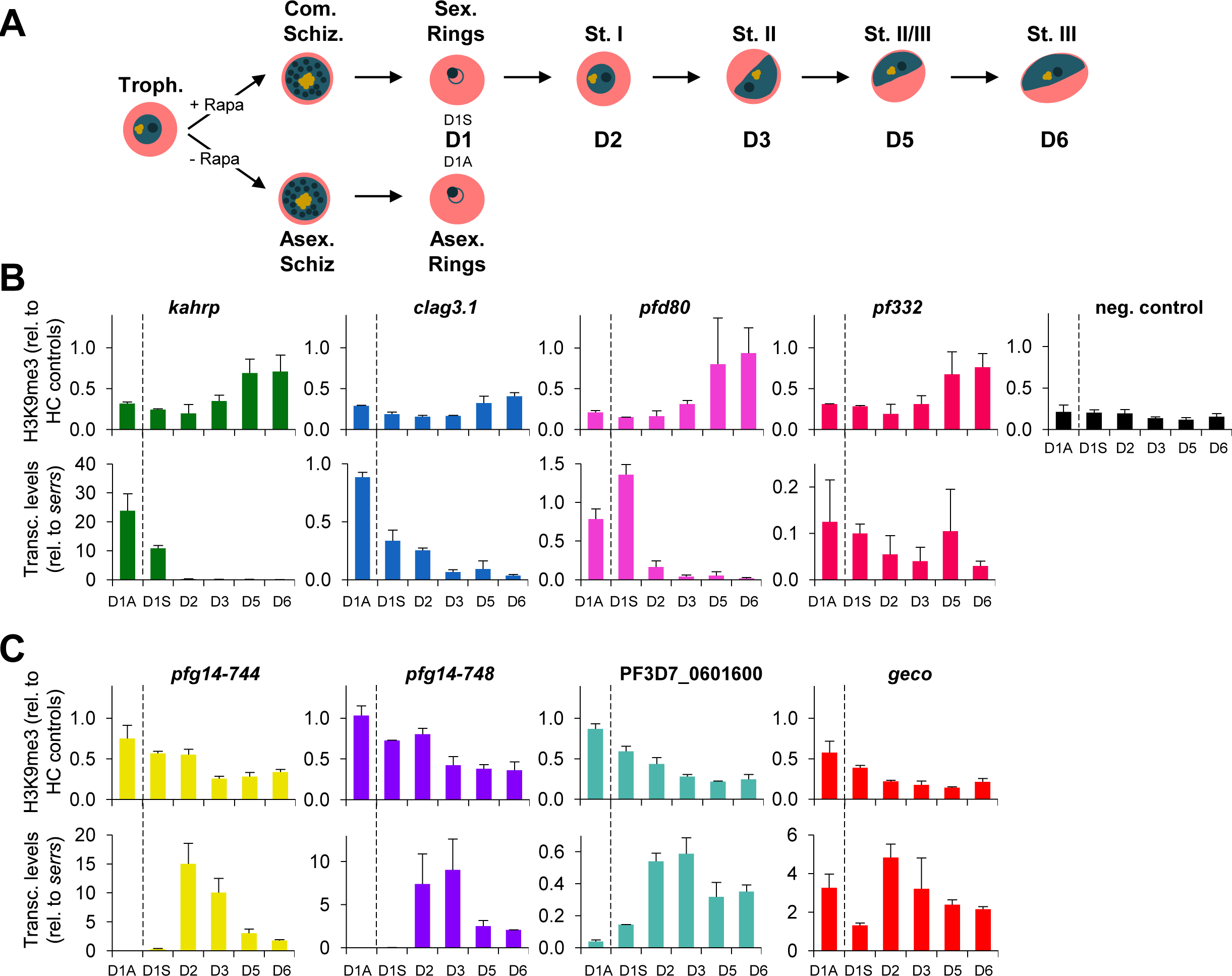
Temporal dynamics of heterochromatin and transcript levels during sexual development. **A.** Schematic of the experiment design. Samples for ChIP-qPCR and RT-qPCR analysis were collected from the same cultures at the days (D) post rapamycin induction indicated. The predominant stage on each day is indicated. Abbreviations of stages are as in Fig. 1. **B-C.** H3K9me3 ChIP-qPCR (top) and RT-qPCR (bottom) analysis for selected genes with increased (**B**) or reduced (**C**) heterochromatin coverage in stage II/III gametocytes in the ChIP-seq experiments. ChIP-qPCR values are recovery (% of input) relative to the average recovery in two heterochromatin-positive control genes (*var* gene PF3D7_1240300 and *clag3.2*). Values for negative controls (euchromatic genes *uce* and *ama1*) are shown at the right. RT-qPCR values are normalised by expression of *serrs* (PF3D7_0717700). All values are the average of two biological replicates, with S.E.M. The vertical dotted line separates asexual from sexual stages.

### Changes in heterochromatin distribution at specific CVG families

Many of the 107 stage II/III differential coverage genes belong to multigene CVG families. Different families of CVGs occupy a relatively conserved position within subtelomeric regions, such that large hypervariable families involved in antigenic variation such as *var*, *rif*, *pfmc-2tm* and *stevor* tend to be located closer to the telomeres, whereas other CVG families that confer functional plasticity to the parasites tend to be located in more telomere-distal positions within subtelomeric regions [39, 40]. Heterochromatin expansion during sexual development mainly affected the latter type of CVG families, in addition to genes that did not belong to gene families previously annotated as containing CVGs. The CVG family with a larger absolute number of genes showing altered heterochromatin occupancy during sexual development was *phist* (23 of 78 genes), involved in erythrocyte remodelling [41–43], followed by *hyp* (11 of 41) that comprises several families involved in the same process [41] and *fikk* (8 of 20), which encodes exported kinases also involved in erythrocyte remodelling [41, 44] (Fig. 5, Sup. Fig. S10 and S11A and Sup. Table S5). Other CVG families with a high proportion (≥ 20%) of differential coverage genes were acyl-coA synthetases (*acs*), acyl-coA binding proteins (*acbp*), glycophorin binding proteins (*gbp*), *clag*, *eba*, *surfin* and *hrp* (Fig. 5, Sup. Fig. S10 and S11A and Sup. Table S5), involved in various processes that include solute transport, erythrocyte invasion and lipid metabolism, among others. As a control, a gene family encoding non-clonally variant essential genes, aminoacyl tRNA synthetases (*ars*), did not show changes in H3K9me3 coverage. Four large families of CVGs located in telomere-proximal positions within subtelomeres (*var*, *rif*, *pfmc2-tm* and *stevor*) showed very few or no heterochromatin changes (Sup. Fig. S10 and S11B).

**Fig. 5.**
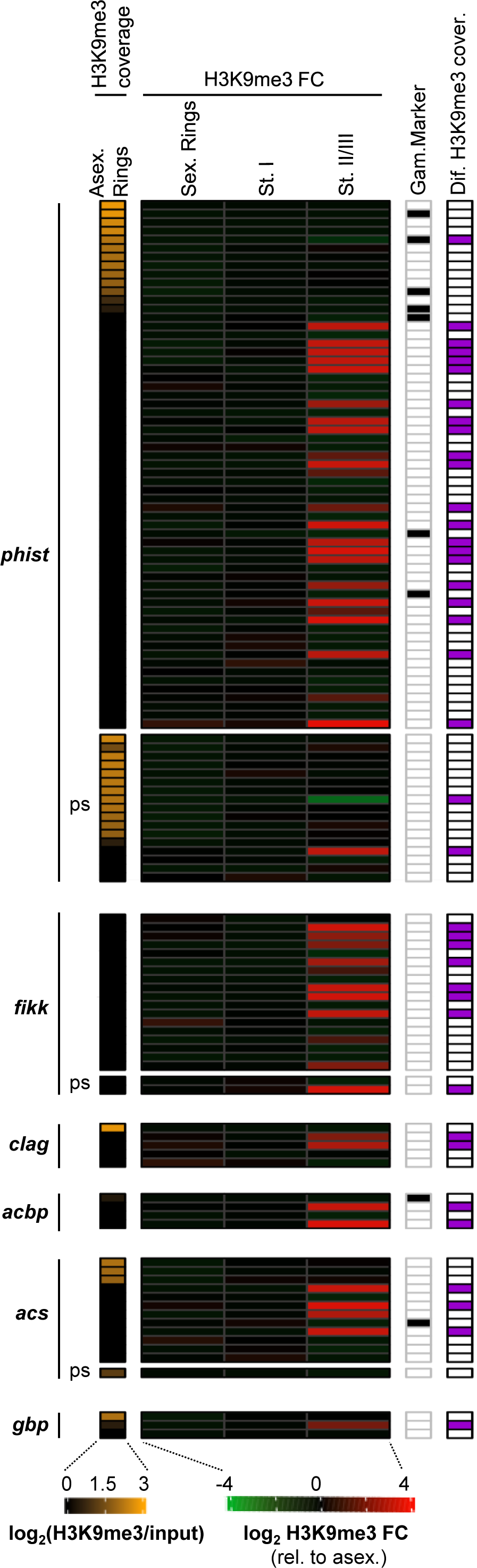
Changes in heterochromatin occupancy during sexual development at specific CVG families. H3K9me3 coverage in E5ind asexual rings (first column) and coverage fold-change (FC) relative to asexual rings at different stages of sexual development. Coverage values were calculated at the region −1,000 to +500 bp from the ATG. Values are the average of two independent biological replicates. All genes of each family are represented, including pseudogenes (ps). The columns at the right indicate whether a gene is a known gametocyte marker, according to a previously published list [9], and whether the gene was classified as a stage II/III differential coverage gene. Abbreviations of stages are as in Fig. 1.

Some gametocyte markers are considered CVGs, as they show variant expression and/or carry heterochromatin marks in asexual parasites. By overlapping previously published lists of CVGs and gametocyte markers [9, 36], we identified 27 genes that fall in this category. It is not known if, once parasites convert into sexual forms, the chromatin state of these genes in asexual parasites affects their expression in gametocytes. In four genes, including *pfg14-744* and *pfg14-748*, H3K9me3 occupancy decreased in putative regulatory regions in stage II/III gametocytes (Sup. Fig. S12 and Sup. Table S6). This result suggests that chromatin remodelling during sexual development secures that these genes are in an euchromatic state in sexually-developing parasites, regardless of their state in asexual parasites. In contrast, *surfin 13.1* and *surfin 4.1* became heterochromatic as a consequence of subtelomeric heterochromatin expansion in stage II/III gametocytes. Of note, *surfin 13.1* has been identified as one of the earliest sexual conversion markers, with high expression already at the sexually-committed schizont stage [19, 45] and reduced expression later during gametocyte development [38]. This raises the intriguing possibility that heterochromatin expansion in stage II/III gametocytes may contribute to developmental silencing of this gene when its product is no longer needed. In the other clonally variant gametocyte genes, including well-known early gametocyte markers (e.g., *gexp5* and *pfg27/25*), H3K9me3 coverage remained invariant at all the stages analysed. Therefore, it remains possible that presence or absence of heterochromatin at these genes in asexual parasites may influence their expression during sexual development, as major heterochromatin changes were not observed at their loci.

### Exploratory analysis to characterise heterochromatin changes at the *pfap2-g* locus after sexual conversion

Our ChIP-seq analysis using the E5ind line does not inform about heterochromatin changes during sexual conversion at the locus encoding the master regulator PfAP2-G, because it has been edited in this transgenic line [19]. The *pfap2-g* locus, which is heterochromatic in asexual parasites [17, 22], is activated by GDV1-mediated heterochromatin displacement in parasites that commit to sexual development [21], but the distribution of heterochromatin at the *pfap2-g* locus soon after sexual conversion has not been characterised.

We used the NF54-*gexp02*-Tom line, in which sexual parasites express a fluorescent marker from the sexual ring stage onwards [46], to sort asexual and sexual rings using flow cytometry and compare their heterochromatin distribution (Sup. Fig. S13A-C). ChIP-seq analysis of the sorted populations revealed reduced heterochromatin occupancy in sexual rings at the region upstream of *pfap2-g*, including the putative promoter, but not at the 5’UTR or coding sequence (Sup. Fig. S13D). This analysis also revealed absence of heterochromatin coverage differences between sexual and asexual rings in genes with differential coverage in sexual rings or stage II/III gametocytes in the E5ind line (Sup. Fig. S13E). This result supports the idea that general heterochromatin changes do not occur until stage II/III gametocytes and the small differences in sexual rings in E5ind were likely attributable to technical reasons. Of note, the analysis of the NF54-*gexp02*-Tom line was complicated by the very small amount of chromatin obtained from sorted parasites. The results of this exploratory analysis will require confirmation using chromatin analysis techniques more suitable for experiments with low input material, such as CUT&TAG or CUT&RUN [47, 48].

## DISCUSSION

Our results show that global heterochromatin changes do not occur at the initial stages of sexual development (i.e., sexual rings and stage I gametocytes) and are only observed from gametocyte stage II/III onwards. Therefore, global heterochromatin redistribution likely plays a role in intermediate or late steps of gametocyte development but not in sexual conversion. The regulatory role of heterochromatin during gametocyte development is additional to its well-established adaptive role in the dynamic regulation of CVGs expression in asexual blood stages [12–14].

Life cycle progression in malaria parasites is mainly regulated by a cascade of ApiAP2 transcriptional regulators, such that expression of different ApiAP2 factors at different stages of the life cycle results in different transcriptomes [49]. For the development of gametocytes, the cascade involves PfAP2-G, PfAP2-G2, PfAP2-G5, AP2-FG and several others [17, 24, 25, 50]. As previously noted [7], the majority of genes that are activated or silenced during gametocyte development do not acquire heterochromatin at any sexual or asexual stage. This indicates that transcription factor networks are the main mechanism governing changes in the transcriptome during gametocyte development, whereas heterochromatin alterations are only a complementary regulatory mechanism. Other changes in chromatin modifications during gametocyte development [5, 51], besides changes in heterochromatin distribution, may also contribute to the regulation of gene expression in gametocytes.

Heterochromatin changes in maturing gametocytes affected several genes previously classified as CVGs [15, 36] and positive for heterochromatin marks in asexual blood stages [6–9], but also a similar number of non-CVGs: several genes that are never heterochromatic during the IDC can be regulated by heterochromatin in gametocytes. Many of these genes belong to multigene families that include other genes that are variantly expressed during the IDC. The majority of the genes with differential heterochromatin coverage in gametocytes encode exported proteins, as previously reported [7], consistent with the prominent role of erythrocyte remodelling during gametocyte development and differences with remodelling during the IDC [52, 53]. Many exported proteins involved in PfEMP1 traffic in asexual blood stages are no longer needed in gametocytes. Some specific gene families such as *phist*, *fikk* or *hyp*, involved in erythrocyte remodelling, are overrepresented among the genes with differential heterochromatin coverage in gametocytes. This may reflect their telomere-distal position within subtelomeric regions, which is the location where heterochromatin expansions occur. It is tempting to speculate that the conserved position of different multigene families within subtelomeric regions [39] may have evolved to facilitate silencing of specific families by heterochromatin spreading during sexual development.

Comparison of our results with previously published data describing the distribution of heterochromatin in sporozoites [30] revealed that the boundaries of subtelomeric heterochromatin domains in sporozoites are more similar to maturing gametocytes than to asexual blood stages. This suggests that the expansion of subtelomeric heterochromatin observed in gametocytes is maintained during development in the mosquito. Previous studies analysing *P. falciparum* blood stages from controlled human malaria infections (CHMI) showed that *var* gene expression patterns are reset during transmission stages [54], and the epigenetic memory for the expression of *clag3* genes [55] and several other (if not all) CVG families is also erased [36]. Furthermore, the observation of heterogeneous CVG expression patterns in blood stage parasites of a volunteer infected by a single sporozoite suggested that the reestablishment of variant gene expression patterns likely occurs during liver stages [36]. Based on these observations and the data presented here, we propose a model for heterochromatin dynamics across the full life cycle (Fig. 6) in which the global heterochromatin distribution in asexual blood stages is first modified in stage II/III gametocytes, involving mainly subtelomeric heterochromatin expansions. After this, heterochromatin distribution is maintained through the rest of gametocyte development and mosquito stages with only small additional changes. A second round of global heterochromatin redistribution occurs during liver stages, before parasites are released again to the bloodstream. This second round of global remodelling reverts subtelomeric heterochromatin expansions and likely involves removal and *de novo* formation of heterochromatin at most loci, resulting in a reset of the expression patterns of specific CVGs. Experiments following changes in heterochromatin distribution in the same parasite line across the full life cycle, which now can be reproduced *in vitro* [56], will be necessary to confirm this model.

**Fig. 6.**
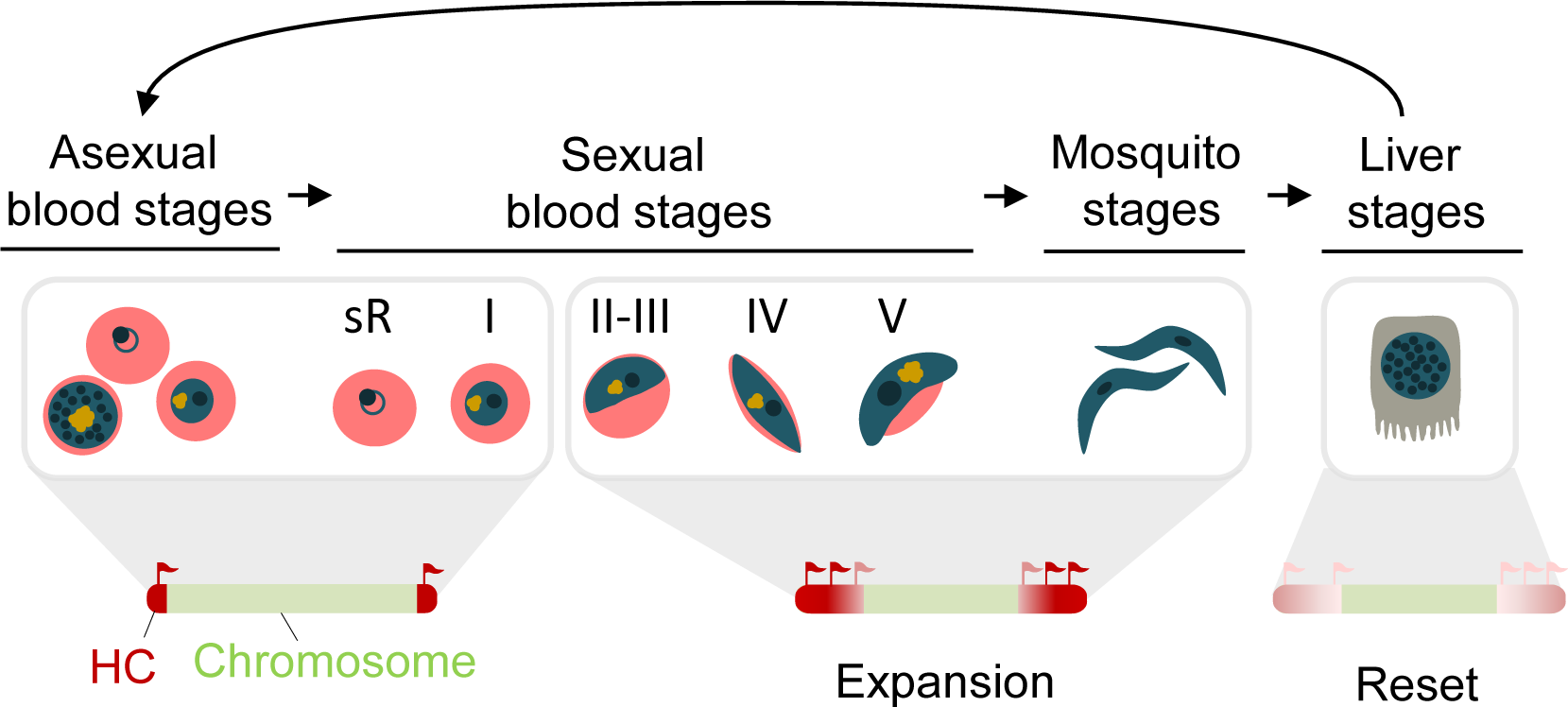
A model for global heterochromatin dynamics across the full life cycle. The extension of subtelomeric heterochromatin domains is maintained throughout all asexual blood stages and early sexual blood stages (sR: sexual rings; I-V: stage I to V gametocytes). From stage II/III gametocytes, there is an expansion of several subtelomeric heterochromatin domains that persists (or even increases) during mosquito stages. In liver stages, there is a hypothetical general remodelling of heterochromatin, such that the epigenetic memory for the transcriptional state of CVGs is erased (reset) and subtelomeric heterochromatin retracts before a new blood infection is established.

The mechanism underlying the expansion of subtelomeric heterochromatin during gametocyte development is not known. Research in model organisms has established that heterochromatin has the ability to spread into adjacent regions in a discontinuous and sequence-independent manner unless limited by boundary elements [1–3], which are not well-described in most eukaryotes, including *Plasmodium* spp. Our data shows that the boundaries for several subtelomeric heterochromatin domains are different between *P. falciparum* asexual blood stages and maturing gametocytes, but conserved between parasite lines of different genetic backgrounds. A possible mechanism that could explain the changes between asexual blood stages and gametocytes involves a different dose of specific heterochromatin regulators between asexual blood stages and gametocytes. The dose of some chromatin modifiers has been shown to determine the extension of heterochromatin domains in *Drosophila* position effect variegation (PEV) and in other systems [1–3, 57]. In this regard, the expression levels of the gene encoding the putative *P. falciparum* H3K9 methyltransferase, PfSet3 (PF3D7_0827800) [8, 58], show a clear increase from roughly stage II/III gametocytes onwards [38], pointing to this protein as a candidate for underlying heterochromatin expansion during gametocyte development.

An unexpected observation from our results was that, in genes with differential heterochromatin coverage between asexual blood stages and gametocytes, transcriptional changes during sexual development preceded changes in heterochromatin occupancy. For genes silenced during sexual development, such as *kahrp* or *pf332*, this may simply reflect that the transcription factors necessary for their expression are already absent before stage II/III gametocytes, as a consequence of the new transcriptional program in gametocytes. For genes activated during sexual development, such as *pfg14-744* and *pfg14-748*, increased transcript levels and also binding of the transcription factor PfAP2-G were first observed in stage I gametocytes, whereas heterochromatin depletion was not observed until stage II/III. This suggests that these genes can be expressed in a heterochromatin environment, which raises the intriguing possibility that PfAP2-G may be a pioneer factor able to bind to and activate its target loci in spite of the presence of heterochromatin. Pioneer factors are a type of transcription factors that can bind DNA wrapped in nucleosomes and in a closed (heterochromatic) chromatin environment, and facilitate transcription by opening chromatin [59]. The possible role of PfAP2-G as a pioneer factor able to activate transcription in a heterochromatic context is suggestive of a different role for heterochromatin in the regulation of CVGs during the IDC or in gametocyte development. In the regulation of CVGs during the IDC, it is well-established that the euchromatic or heterochromatic state at regulatory regions is associated with active transcription or silencing, respectively, indicating that heterochromatin actually determines gene silencing [9, 26–28]. Additionally, the euchromatic or heterochromatic state is conserved through stages of the IDC at which the gene is not expressed, maintaining the epigenetic memory [12]. In contrast, the role of heterochromatin changes in gametocyte development may be less deterministic and simply contribute to lock down the active or silenced transcriptional states.

## MATERIALS & METHODS

### Parasite cultures and preparation of sexual and asexual parasite samples

The E5ind and NF54-*gexp02*-Tom lines have been previously described [19, 46]. Cultures were regularly maintained under standard *P. falciparum* culture medium containing Albumax II and no human serum. The E5ind line was cultured under 10 nM WR99210 (Jacobus Pharmaceuticals Co., USA) pressure to maintain the *cam* promoter integrated at the *pfap2-g* locus in an active state, but drug pressure was removed at the cycle of induction [19]. The NF54-*gexp02*-Tom line was regularly maintained in medium supplemented with 2 mM choline (Sigma-Aldrich, C7527) to repress sexual conversion [46].

Induction of sexual conversion in the E5ind line was performed as previously described by adding 10 nM rapamycin (Sigma-Aldrich, R0395) for 1 h to sorbitol-synchronised cultures at the trophozoite stage (20 h post-synchronisation) [19]. Control cultures were treated, in parallel, with DMSO solvent. From one cycle before induction, cultures were maintained in erythrocytes filtered with Plasmodipur columns (R-Biopharm, 8011FILTER10U) to remove residual leukocytes and avoid human DNA contamination. After reinvasion (about 24 h after induction), 2 mM choline and 50 mM N-acetylglucosamine (Sigma-Aldrich, A3286) were added to the cultures, to improve gametocyte maturation and to prevent asexual parasite growth, respectively. Recombination at the edited *pfap2-g* locus after induction was confirmed by PCR analysis of genomic DNA extracted ∼24 h after induction, as described [19].

To obtain TdTomato positive (TdTom+) and negative (TdTom-) rings, NF54-*gexp02*-Tom cultures were tightly synchronised to a 5 h age window using Percoll purification of mature forms followed by sorbitol lysis to eliminate mature forms 5 h later [60]. Immediately after sorbitol lysis, when parasites were 0-5 h post-invasion (hpi), the choline supplement was removed to stimulate sexual conversion [21, 61, 62]. The sexual conversion rate, determined by flow cytometry [46], was ∼50% in this experiment. After reinvasion, at ∼11-16 hpi of the next cycle, TdTom+ and TdTom-parasites were sorted in a BD FACS Aria SORP cell sorter. Erythrocytes were gated on SSC-A/FSC-A and then single cells were gated on FSC-H/FSC-A to avoid cell doublets or aggregates. Next, TdTom+ and TdTom-cells were separated using the combination of 488 nm and 561 nm lasers (Sup. Fig. S13B). Sorting parameters included: sheath pressure set at 20 psi, use of a 100 µm nozzle tip and temperature maintained at 37°C. Sorted cells were collected into tubes containing culture medium supplemented with 2 mM choline. We collected a total of 3.87×10^6^ TdTom+ cells (sexual rings) and 7.8×10^7^ TdTom-cells (containing 3.74×10^6^ asexual rings and uninfected erythrocytes). Sorting took approximately 7 h. The purity of the resultant TdTom+ and TdTom-preparations was >90%, according to flow cytometry analysis (Sup. Fig. S13C). Uninfected erythrocytes were added to the TdTom+ sorted cells to adjust the parasitaemia to that of the TdTom-sorted cells (∼5%).

### ChIP-seq experiments

ChIP-seq experiments were performed essentially as described [19, 36]. Parasites were isolated from erythrocytes using saponin lysis. After formaldehyde cross-linking, chromatin was extracted with the MAGnify kit (Life Technologies). Sonication to obtain 150 - 200 bp DNA fragments was performed with a Covaris M220 apparatus. Immunoprecipitation was done with 2 µg of chromatin and 4 µg of antibody against H3K9me3 (Diagenode, C15410193, lot A2217P). Sequencing libraries were prepared from 5 ng of immunoprecipitated DNA as previously described [19, 36], using conditions adapted to the extremely AT-rich *P. falciparum* genome [63]. NEBNext Multiplex Oligos for Illumina (New England BioLabs) were used for multiplexing, and 8 to 10 PCR cycles with the KAPA HiFi PCR kit (Kapa Biosystems) were used for amplification. For all purification steps, AMPure XP beads (Beckman Coulter, A63882) were used. Sequencing was performed using an Illumina NextSeq 550 system, obtaining 8 to 15 million 150 bp paired-end reads per sample.

For the ChIP-seq analysis of NF54-*gexp02*-Tom FACS-sorted parasites, we used a similar protocol but library preparation was performed using the NEBNext Ultra DNA Library Prep Kit (New England BioLabs, ref. E7370) for Illumina following manufacturer’s specifications except for end repair, which was performed for 1 h at 45°C (instead of 30 min at 65°C). DNA was amplified for 15-19 cycles.

### RT-qPCR and ChIP-qPCR assays

ChIP-qPCR analysis of uninduced E5ind cultures at the asexual ring stage and induced cultures at different sexual stages was performed approximately as described [64]. In brief, chromatin was extracted as for the ChIP-seq experiments, but DNA was sonicated using a Bioruptor 300 (Diagenode) to obtain ∼500 bp fragments. After immunoprecipitation (as for ChIP-seq experiments), input and immunoprecipitated samples were analysed by qPCR in triplicate wells using the standard curve method (with a standard curve included in each plate for each primer pair) and the primers described in Supplementary Table S8. In each experiment, we included *ama1* and *uce* as heterochromatin negative controls and the *var* gene PF3D7_1240300 and *clag3.2* as heterochromatin positive controls.

RNA for RT-qPCR analysis was prepared using the TRIzol method, treated with DNAse I and purified as previously described [60, 64]. Reverse transcription with a mixture of random and oligo (dT) primers and qPCR analysis of the cDNAs using the PowerSYBR Green master mix (Applied Biosystems) and the standard curve method were also performed as previously described [60, 64]. All primers used are listed in Sup. Table S8. Transcript levels were normalised against the serine–tRNA ligase gene (*serrs*, PF3D7_0717700), which is expressed at relatively stable levels in all blood stages except for very young rings, which were not included in this study.

### ChIP-seq data analysis

New ChIP-seq data generated in this study as well as previously published data [7, 30] were analysed approximately as previously described [9]. After quality control and trimming, reads were aligned against the *P. falciparum* 3D7 genome (PlasmoDB v. 55) using bowtie2 (v. 2.3.0). Duplicates reads were removed and peak calling for each individual sample was performed using MACS2 software (v. 2.2.7) with parameters: -f BAMPE -B -g 2.41e7 --keep-dup all --fe-cutoff 1.5 -nomodel --extsize 150. IP tracks were normalised against their inputs using the Deeptools suite, with the bamCompare option. Coverage at specific genome regions was calculated from normalised mean log_2_ (IP/input) bedgraph tracks, using the bedtools -map option.

To calculate the proportion of the genome of each parasite line covered by a MACS2 H3K9me3 ChIP-seq speak, each .narrowPeak file containing all peak intervals were converted into a fasta file using Bedtools (-getfasta option) and, after counting the number of base pairs covered by the mark, it was divided by the size of the full genome.

To identify differential peaks, which here refers to regions with differential coverage between asexual rings and sexual stages, we used a previously described in-house method [9], with some modifications. Each sexual stage was compared with asexual rings from the same experiment. In brief, we first calculated the input-normalised coverage for all 100 bp bins overlapping a MACS2 peak in one or more of the samples and adjusted the distribution of coverage intensities to a gaussian mixture distribution with two components (the rationale being that one component accounts for the non-enriched regions and the other one for the enriched regions). Next, we calculated for each bin in each sample a cumulative density function (CDF) value for each of the two components. Bins with a CDF value difference of >0.2 (for any of the two components) were classified as having differential coverage. We merged bins with differential coverage separated by <1,000 bp, and filtered out differential peaks of <150 bp after merging. Finally, we only retained as differential peaks the intervals that were common between the two replicates and removed regions of ≤500 bp or with an absolute value of the log_2_ of the coverage fold-change between the sexual stage and asexual rings of <1. The final differential peaks were then intersected with protein coding genes (either the −1,000 to +500 bp from the ATG region or the rest of the annotated transcript products, including 5’UTR and 3’UTR when available in PlasmoDB) and other genetic elements. Genes for which the −1,000 to +500 bp region overlapped with a differential coverage peak were retained in the differential coverage gene lists if the log_2_ of the H3K9me3 coverage fold-change between asexual rings and the sexual stage tested was ≥1 in this −1,000 to +500 bp region.

### Determination of the extension of heterochromatin domains at subtelomeric regions or internal islands

From the initial .narrowPeak files generated by individual MACS2 peak calling, we merged peaks separated by ≤ 10 Kb using bedtools (parameters merge -i .narrowPeak -d 10000) and eliminated the peaks with a length <500 bp. Next, to define the full extent of each heterochromatin domain we merged regions separated by ≤60 Kb and, for each subtelomeric region, extended the regions to the extreme of the chromosome regardless of whether or not it was covered by a MACS2 peak (Sup. Fig. S3A). Finally, we used visual inspection of IGV tracks to confirm consistency between replicates and edited manually the limits of the heterochromatin region in one of the replicates when there was a large discrepancy with the other replicate and an error of the algorithm was apparent (one subtelomeric domain was manually edited).

## ACKNOWLEDGMENTS

We are grateful to Alba Pérez-Cantero for help with training and advice with parasite cultures and ChIP experiments, and for critical reading of the manuscript. This work was supported by grants PID2019-107232RB-I00 and PID2022-137863OB-I00 to A.C. from the Spanish Ministry of Science and Innovation (MCI)/ Agencia Estatal de Investigación (AEI, 10.13039/501100011033), co-funded by the European Regional Development Fund (ERDF, European Union), and “la Caixa” Banking Foundation (HR18-00267 to A.C.). L.M.-T. was supported by a fellowship from the Spanish Ministry of Economy and Competitiveness (BES-2017-081079), co-funded by the European Social Fund (ESF). Our research is part of ISGlobal’s Program on the Molecular Mechanisms of Malaria, which is partially supported by the Fundación Ramón Areces. We acknowledge support from the Spanish Ministry of Science and Innovation through the “Centro de Excelencia Severo Ochoa 2019-2023” Program (CEX2018-000806-S), and support from the Generalitat de Catalunya through the CERCA Program.

## DATA AVAILABILITY

The new ChIP-seq data presented in this article was deposited in the GEO database with accession n. GSE252334. ChIP-seq data can also be visualised at the UCSC genome browser (https://genome.ucsc.edu/cgi-bin/hgTracks?db=hub_2790993_GCA_000002765.3&lastVirtModeType=default&lastVirtModeExtraState=&virtModeType=default&virtMode=0&nonVirtPosition=&position=Pf3D7_07_v3%3A1%2D206072&hgsid=2017847446_TqNzA9wD60Du0pJmFtzU0ySlAQav).

